# Prefrontal and Parietal Attractor Networks Mediate Working Memory Judgments

**DOI:** 10.1101/2020.06.01.128173

**Authors:** Sihai Li, Christos Constantinidis, Xue-Lian Qi

## Abstract

The dorsolateral prefrontal cortex plays a critical role in spatial working memory and its activity predicts behavioral responses in delayed response tasks. Here we addressed whether this predictive ability extends to categorical judgments based on information retained in working memory, and is present in other brain areas. We trained monkeys in a novel, Match-Stay, Nonmatch-Go task, which required them to observe two stimuli presented in sequence with an intervening delay period between them. If the two stimuli were different, the monkeys had to saccade to the location of the second stimulus; if they were the same, they held fixation. Neurophysiological recordings were performed in areas 8a and 46 of the dlPFC and 7a and lateral intraparietal cortex (LIP) of the PPC. We hypothesized that random drifts causing the peak activity of the network to move away from the first stimulus location and towards the location of the second stimulus would result in categorical errors. Indeed, for both areas, when the first stimulus appeared in a neuron’s preferred location, the neuron showed significantly higher firing rates in correct than in error trials. When the first stimulus appeared at a nonpreferred location and the second stimulus at a preferred, activity in error trials was higher than in correct. The results indicate that the activity of both dlPFC and PPC neurons is predictive of categorical judgments of information maintained in working memory, and the magnitude of neuronal firing rate deviations is revealing of the contents of working memory as it determines performance.

**SIGNIFICANCE STATEMENT:** The neural basis of working memory and the areas mediating this function is a topic of controversy. Persistent activity in the prefrontal cortex has traditionally been thought to be the neural correlate of working memory, however recent studies have proposed alternative mechanisms and brain areas. Here we show that persistent activity in both the dorsolateral prefrontal cortex and posterior parietal cortex predicts behavior in a working memory task that requires a categorical judgement. Our results offer support to the idea that a network of neurons in both areas act as an attractor network that maintains information in working memory, which informs behavior.

## INTRODUCTION

Working memory, the ability to maintain and manipulate information over a time span of seconds, is a core component of higher cognitive functions (Baddeley, 2012). Neurophysiological experiments in the prefrontal cortex of non-human primates identified neurons that remain active during a period of seconds over which a stimulus is maintained in memory (Fuster and Alexander, 1971; Funahashi et al., 1989). This “persistent activity” is thought to be maintained through recurrent connections in a network of neurons (Chaudhuri and Fiete, 2016; Zylberberg and Strowbridge, 2017). Individual neurons exhibiting persistent activity are selective for the properties of stimuli held in memory (Qi and Constantinidis, 2013) and trials in which persistent activity is diminished are more likely to result in errors (Funahashi et al., 1989; Zhou et al., 2013). It is therefore believed that persistent discharges constitute the neural correlate of working memory (Constantinidis and Klingberg, 2016).

In recent years, the neural mechanisms of working memory have come under debate (Constantinidis et al., 2018; Miller et al., 2018). Short-term synaptic changes and rhythmic bursts in the gamma frequency range have been proposed as the main mechanisms of working memory generation (Stokes, 2015; Lundqvist et al., 2016). Perhaps the strongest argument in favor of the persistent discharge model, is that variations in persistent activity during the delay interval of working memory tasks are strongly predictive of behavior (Riley and Constantinidis, 2016). For example, persistent activity recorded from trials in which monkeys made eye movements deviating clockwise vs. counterclockwise relative to the true location of the stimulus yields slightly different tuning curves, as would be expected if the location recalled was determined by the peak of activity at the end of the delay period in a bump attractor network (Wimmer et al., 2014; Barbosa et al., 2019). Nonetheless, such results may be interpreted as suggestive of motor preparation rather than working memory itself (Lundqvist et al., 2018).

At the same time, the site of memory maintenance has come under debate. fMRI studies have been successful in decoding information held in memory from the visual cortex (Harrison and Tong, 2009; Albers et al., 2013; Ester et al., 2013; Xing et al., 2013; Sreenivasan et al., 2014a). Visual cortical areas rather than the prefrontal cortex have been thus suggested as the site of information maintenance (Sreenivasan et al., 2014b; Christophel et al., 2017). Activation of the posterior parietal cortex in fMRI studies have also shown to best predict individual working memory capacity (Todd and Marois, 2004, 2005).

We were thus motivated to test two basic tenets of current working memory models. First, we devised a novel working memory task, allowing us to determine if the variability of persistent discharges in the prefrontal cortex is predictive of working memory behavior in a task that dissociates stimulus maintenance from motor preparation. Secondly, we wished to ascertain if such a relationship is exclusive to the prefrontal cortex, or is also present in a more posterior cortical region implicated in spatial working memory, the posterior parietal cortex (Constantinidis et al., 2013).

## METHODS

Two male rhesus monkeys (*Macaca mulatta*) weighing 9-12 kg were used in these experiments. Neural recordings were carried out in areas 8 and 46 of the dorsolateral prefrontal cortex and areas 7a and lateral intraparietal area (LIP) of the posterior parietal cortex. All experimental procedures followed guidelines by the U.S. Public Health Service Policy on Humane Care and Use of Laboratory Animals and the National Research Council’s Guide for the Care and Use of Laboratory Animals and were reviewed and approved by the Wake Forest University Institutional Animal Care and Use Committee.

### Experimental Setup

Monkeys sat in a primate chair with their head fixed while viewing an LCD monitor positioned 68 cm away from their eyes with dim ambient illumination. Animals were required to fixate a 0.2° white square appearing in the center of the monitor screen. During each trial, animals had to maintain fixation on the square while visual stimuli were presented either at a peripheral location or over the fovea in order to receive a liquid reward. Any break of fixation immediately terminated the trial and no reward was given. Eye position was monitored throughout the trial using a non-invasive, infra-red eye position scanning system (model RK-716; ISCAN, Burlington, MA). The system achieved a <0.3° resolution around the center of vision. Eye position was sampled at 240 Hz, digitized and recorded. Visual stimuli display, monitoring of eye position, and the synchronization of stimuli with neurophysiological data were performed with in-house software (Meyer and Constantinidis, 2005) implemented on the MATLAB environment (Mathworks, Natick, MA).

### Task and Stimuli

Monkeys were trained to perform the Oculomotor Delayed Response task (ODR, Fig. 1A-B) and the Match-Stay Nonmatch-Go (MSNG) task (Fig.1C-D). During recording days, the session started with an ODR task, which allowed us to determine the presence of neurons which responded during the delay period and the approximate location of their receptive fields. The MSNG task followed. Based on estimated best neuronal responding location, we determined the locations of stimuli used in the MSNG task, however recordings were obtained from neurons at multiple electrodes and the stimulus location could appear anywhere relative to a neuron’s receptive field.

**Figure 1.**
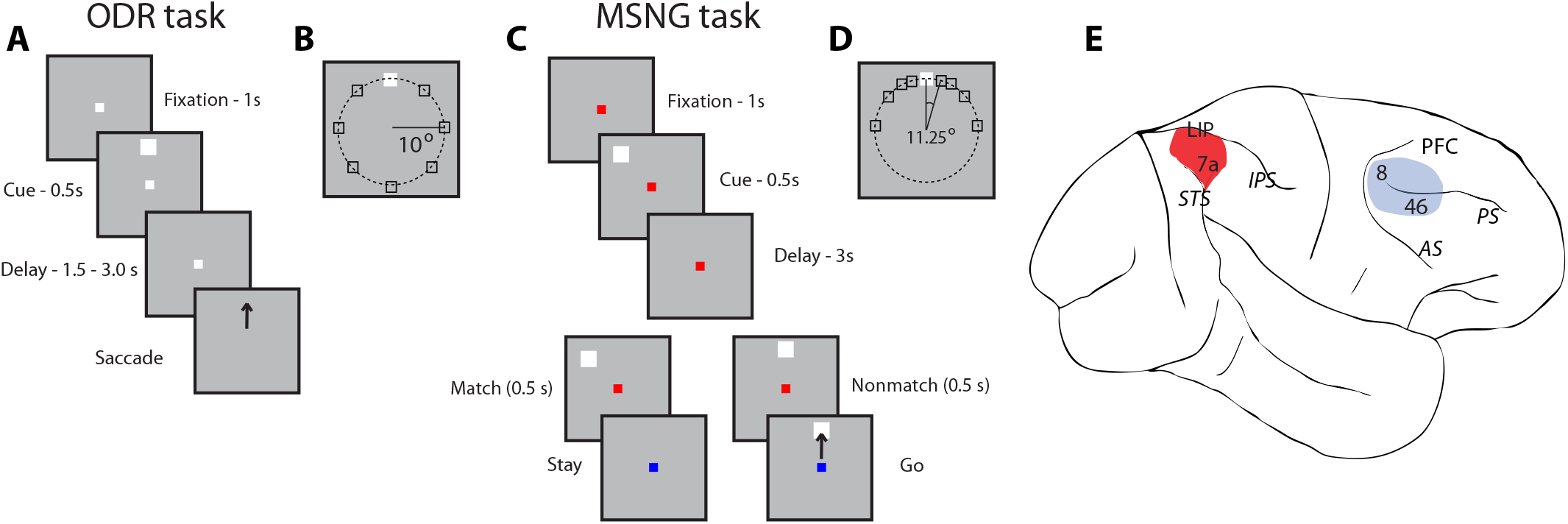
Tasks and areas for neurophysiological recordings. (A) Frames represent the sequence of events during the ODR task. The monkey is required to observe a cue stimulus, maintain fixation during delay period, and make eye movement to the remembered location of the visual cue once the fixation point disappears. (B) Possible locations of visual stimuli on the screen in the ODR task. (C) Sequence of events in the MSNG task. The monkey is required to observe the first cue and maintain fixation during the delay period. Then another visual stimulus appears, and a monkey needs to determine if it appeared at the same location as the cue. If two visual stimuli appeared at the same location (they match), the monkey needs to stay at the fixation point, after its color changes. If the location of the second stimulus deviates from the first (nonmatch), the monkey is required to make an eye movement to the (visible) second stimulus once the color of fixation point switches. (D) Possible locations of visual stimuli presentation on the screen in the MSNG task. White square represents the reference stimulus, around which all other stimuli are presented in a daily session. The reference stimulus appeared at different locations in different sessions, typically at the flank of the neuron’s tuning curve. (E) Regions of neurophysiological recordings, including areas 8 and 46 in dlPFC (highlighted in blue) and areas 7a and LIP in PPC (highlighted in red). Abbreviations: AS, Arcuate Sulcus; PS, Principal Sulcus; IPS, Intraparietal Sulcus; STS, Superior Temporal Sulcus.

The ODR task required monkeys to remember the spatial location of a 1° white cue stimulus displayed on a screen for 0.5s and after a delay period of either 1.5 s or 3s, to make a saccade to its remembered location (Fig. 1A). Stimuli could appear at any of eight locations arranged on an (invisible) circle of 10 degrees of visual angle eccentricity (Fig. 1B), and the monkeys were required to make an eye movement to the remembered location of the cue within 0.6s to receive a liquid reward. Correct responses were considered those in which the saccadic end point deviated no more than 5-6° from the center of the stimulus (3-4° from the edge of the stimulus), and the monkey held fixation within this window for 0.1s.

The MSNG task required the monkeys to remember the location of the cue presented in the same fashion, but after a 3s delay period, a second stimulus appeared, either at the identical location (match) or a different location (nonmatch, Fig. 1C). After 500 ms, the fixation point changed color, and if the second stimulus was a match, the monkey was required to maintain fixation; if the second stimulus was a nonmatch, the monkey was required to make a saccade towards this visible stimulus. The monkeys received a liquid reward for a correct response. In each daily session, the cue could appear pseudo-randomly at one of nine possible locations (Fig. 1D). Possible cue locations included a reference location (white square in Fig. 1D) and eight locations deviating from the reference location by an angular distance of 11.25°, 22.5°, 45° and 90°, clockwise and counterclockwise. The cue was followed by a matching stimulus appearing at the identical location in approximately half the trials (9/17 conditions) and by a nonmatch stimulus always appearing at the reference location in the remaining trials (8/17 conditions). The reference location varied from session to session. During the course of the recording session, we tried to select a reference location that appeared at the flank of the tuning curve of neurons under study, based on their responses in the ODR task, which preceded the MSNG task. However, multiple neurons were recorded in each session and the location of the reference stimulus could not be optimized for all at the same time. Therefore, in practice, the reference location could appear at any position relative to the neuron’s tuning curve and this determination was made post hoc. All stimuli were presented at an eccentricity of 10 degrees of visual angle, as in the ODR task. During training, other relative locations of cue and nonmatch stimuli were also displayed, to ensure that the monkey understood the rule and could perform the task for any combination of stimuli.

### Surgery and neurophysiology

Two, 20-mm diameter craniotomies were performed over the lateral prefrontal cortex and the posterior parietal cortex and a recording cylinder was implanted over each site’s right hemisphere, respectively. The location of the cylinder was visualized with anatomical MRI imaging and stereotaxic coordinates post-surgery. Neurophysiological recordings were obtained as we have described before (Zhou et al., 2016b). Briefly, we used tungsten-coated electrodes with a 200 or 250 μm diameter and 4 MΩ impedance at 1 kHz (FHC, Bowdoinham, ME). Arrays of up to 4-microelectrodes spaced 0.5-1 mm apart were advanced into the cortex with a Microdrive system (EPS drive, Alpha-Omega Engineering) through the dura into the cortex. The signal from each electrode was amplified and band-pass filtered between 500 Hz and 8 kHz while being recorded with a modular data acquisition system (APM system, FHC, Bowdoin, ME). Waveforms that exceeded a user-defined threshold were sampled at 25 μs resolution, digitized, and stored for off-line analysis.

### Anatomical Localization

Neural recordings were performed in two cortical areas, the dorsolateral prefrontal cortex and the posterior parietal cortex (Fig 1E). Prefrontal recordings included areas 46 and 8 containing the caudal part of both banks of the Principal Sulcus, and the area between the Principal sulcus and the Arcuate sulcus. Posterior parietal recordings encompassed area 7a and LIP, which is directly connected to the dorsolateral prefrontal cortex (Cavada and Goldman-Rakic, 1989). During the experimental sessions, the travel distance of electrodes before entering the cortex allowed us to map the sulcal pattern of each cylinder and provided a coarse map of anatomical location. Upon completion of the experiments, the anatomical location of electrode penetration was recorded onto an image of cortical surface obtained through previous MRI scanning.

### Behavioral Data Analysis

All analysis of behavioral (and neural) data was performed in the MATLAB environment (Mathworks, Natick, MA, version 2012a-2019). We expressed the correct performance in the ODR task and the MSNG task as the percentage of trials that resulted in correct responses. Some trials were aborted early due to breaks in fixation, blinks, or premature saccades, before the fixation point changed color. These trials were ignored in performance estimation. We additionally calculated d’ (sensitivity index) defined as d’ = Z(hit rate) – Z(false alarm rate) where Z(x) is the inverse of the cumulative distribution function of the Gaussian distribution. The d’ value was calculated on a session by session basis, based on the MATLAB norminv function. Hit rate in this context represents the percentage of correct detections of nonmatch trials, and false alarm the error rate in match trials that the monkey incorrectly perceived as Nonmatch trial and made a saccade towards the second stimulus.

### Neural Data Analysis

Recorded spike waveforms were sorted into separate units using an automated cluster analysis relying on the KlustaKwik algorithm (Harris et al., 2000), which applied principal component analysis of the waveforms. Mean firing rate was then determined in each trial epoch. To ensure the stability of firing rate in the recordings analyzed, we identified recordings in which a significant effect of trial sequence was evident on the baseline firing rate (ANOVA, p<0.05), e.g. due to a neuron disappearing or appearing during a run, as we were collecting data from multiple electrodes. Data from these sessions were truncated so that analysis was only performed on a range of trials with stable firing rate.

We identified task-related neurons as those with firing rates during the first stimulus presentation or delay period that were higher compared to the 1s fixation period preceding it, based on a paired t-test, evaluated at the p<0.05 level. Population discharge rates were evaluated by averaging activity from multiple neurons and constructing Peri-Stimulus Time Histograms (PSTH). These were constructed using the best stimulus responses for each neuron. PSTHs were aligned to the cue presentation and averaged responses for each stimulus set and brain region.

Correct and error conditions were compared for nonmatch trials in which the cue stimulus appeared at a location that corresponded to a higher firing rate than the second stimulus location based on the neuron’s tuning curve (preferred cue). As mentioned in the description of the MSNG task, above, the location of the cue (and match) varied across conditions, whereas the location of the nonmatch stimulus was fixed. It was possible, therefore, to identify trials in which the cue appeared at a more preferred location than the nonmatch location. Firing rate was also compared for trials in which the cue stimulus appeared at a location that corresponded to a lower firing rate than the second stimulus location (non-preferred cue). This determination was done separately for each neuron. A minimum of 2 error trials was required for a neuron to be included in this comparison and not all neurons were used for all conditions, resulting in a slightly different sample size for each test.

In order to quantify the trial-to-trial association between perceptual choice and neuronal activity, we analyzed trials that resulted in correct choices and incorrect choices in the delayed match-to-sample task and the reaction-time task using the choice probability analysis based on signal detection theory (Britten et al., 1996a). Firing rates of trials involving the same cue and nonmatch stimulus were pooled separately for correct and error outcomes. A receiver operating characteristic (ROC) curve was computed from these two distributions of firing rates. The choice probability, a measurement of correlation between the behavioral choice and neuronal activity, was defined as the area under the ROC curve. A choice probability value of 1 indicates that there is a perfect correlation between the behavioral choices and the neuronal discharge rates; a value of 0.5 indicates no correlation between the two. Time-resolved choice probabilities were computed from the spikes in 250 ms time windows, stepped by 50 ms intervals. This analysis was done separately for the preferred and non-preferred cue conditions. Results from all available neurons were averaged together to produce population responses.

## RESULTS

### Behavioral Performance

Two monkeys were trained to perform the Oculomotor Delayed Response (ODR) and a novel, Match-Stay, Nonmatch-Go (MSNG) task (Fig. 1). The MSNG task required monkeys to observe and remember a visual cue and, after a 3s delay period, observe a second cue and compare its location to the remembered location of the first visual cue. If the two stimuli were displayed at the same location, they defined a Match trial and the monkey should hold fixation (stay). If the locations of two visual cues differed from each other, they defined a Nonmatch trial and the monkey was required to make an eye movement to the second stimulus, which remained visible at the screen at that point (go). The monkeys were rewarded for all correct stay or go responses. Trials were aborted immediately if the monkey made a premature eye movement. Stimuli were displayed at 9 possible locations, deviating from a reference cue (shown in white in Fig. 1D) by 11.25°, 22.5°, 45° or 90°, clockwise and counterclockwise. Match trials involved successive presentation of two stimuli at any of these 9 locations. Nonmatch trials involved presentation of the first stimulus at one of the eight locations around the reference location, and the second stimulus always at the reference location. The reference location changed from session to session. During training, the monkeys were exposed to other combinations of match and nonmatch location so that they mastered the rule regardless of first and second stimulus location.

Recordings commenced once each monkey reached asymptotic performance and no further consistent improvement was observed. The two monkeys (KE and LE) performed a total of 43078 complete (non-aborted) trials during recording sessions. In Nonmatch trials, performance increased monotonically as a function of increasing angular distance between the first and second stimulus (Fig 2A-B). In other words, the further away the second stimulus was from the first, the more likely that the animal perceived the location of the second stimulus as different from the remembered location of the first stimulus. For stimuli separated by 11.25°, 22.5°, 45° and 90°, mean performance across daily sessions for monkey KE was 62%, 71%, 86% and 87%, respectively (Fig. 2A); for monkey LE 54%, 66%, 78% and 85%, respectively (Fig. 2B). Performance in match trials was intermediate (KE, 82%; LE 79% correct responses). Performance was also expressed as a sensitivity index (d’) which provides the separation between means of the signal and the noise distribution, in this case the correct detections of nonmatch and false alarms of match trials perceived as nonmatch. A monotonic increase in sensitivity index was observed with larger angular distance between two stimuli. For monkey KE that was 1.27, 1.56, 2.03 and 2.12, for the first 4 separations respectively (Fig. 2C); for monkey LE, 0.92, 1.24, 1.62, and 1.84, respectively (Fig. 2D).

**Figure 2.**
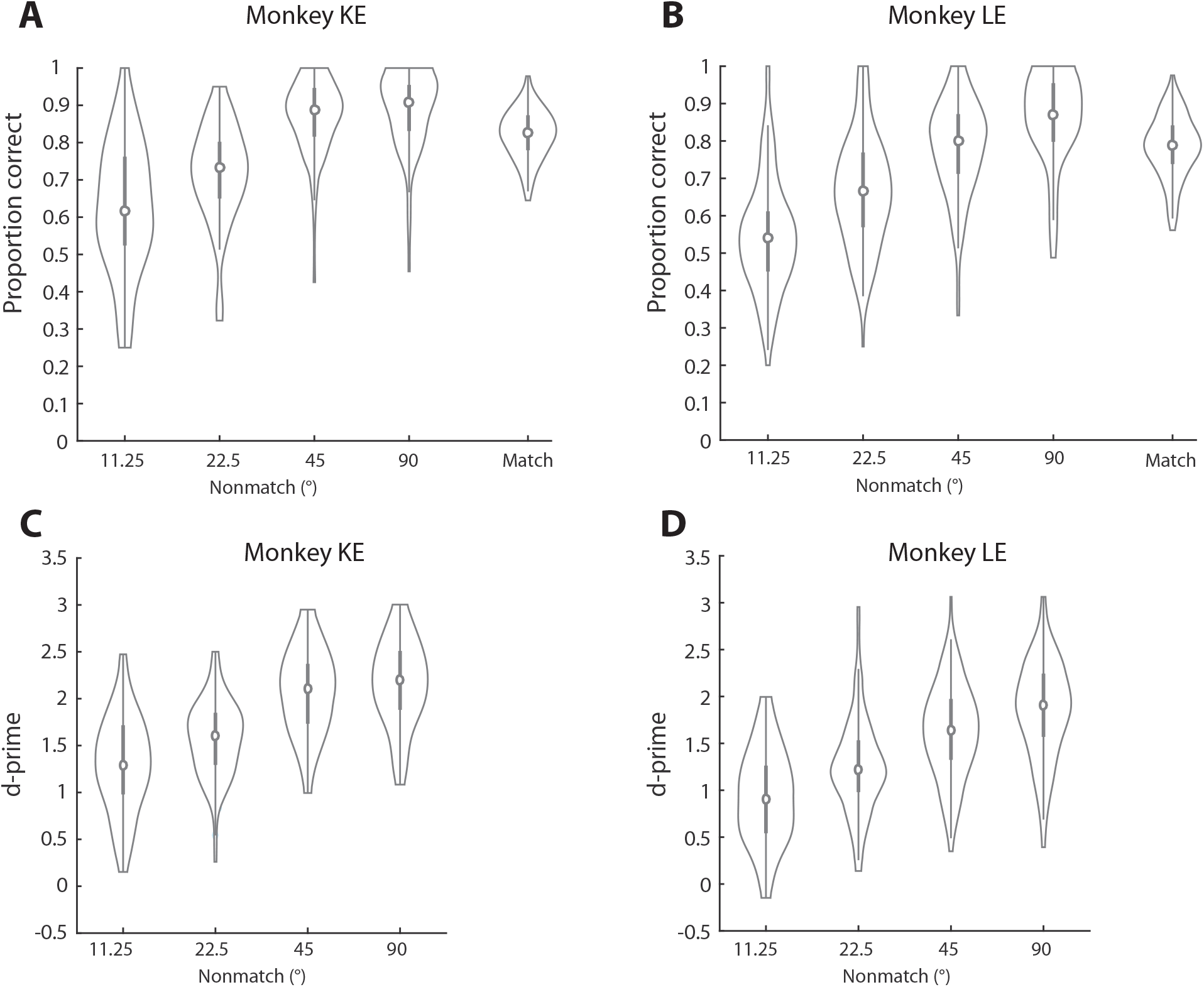
Behavioral performance. (A) Performance of monkey KE in the MSNG task, during neural recording sessions (n = 82). Proportions of correct trials for Nonmatch trials are plotted as a function of the angular distance between the two consecutive stimuli. Performance in Match trials is also plotted, for comparison. The estimate of performance excludes trials aborted before the color of fixation switched due to breaks in fixation. Each colored dot represents the proportion of correct trials in a single session. The median (circle) and interquartile difference (gray vertical line) are also indicated in the plot. (B) Behavioral performance for monkey LE in the MSNG task (n = 132) plotted as in A. (C) Sensitivity index (d-prime) for monkey KE, calculated based on the correct detections of nonmatch stimuli and false reports of match stimuli as nonmatches. Sensitivity measures are plotted for different angular distances between the two consecutive stimuli. Each colored dot represents the d-prime value calculated in a single session. (D) As in C, sensitivity index for monkey LE in the MSNG task.

### Task-related neuronal responses

Neuronal activity was recorded from areas 8 and 46 of the dorsolateral prefrontal cortex and areas 7a and LIP of the posterior parietal cortex. Responses were recorded from 1378 neurons in two cortical areas from two monkeys as the monkeys performed MSNG task (monkey KE – 598 neurons, LE – 780 neurons). Among these neurons, 144 PFC neurons and 145 PPC neurons exhibited significantly elevated responses in the first cue period or the delay period, compared to the baseline activity (paired *t*-test, P < 0.05) and were selected for further analysis. Neurons in both brain regions generated activity during the ODR and MSNG tasks. The population responses are shown in Figure 3. Overall, the absolute firing rate was slightly higher in the prefrontal cortex in our sample, however, both regions generated persistent activity (Fig. 3). Responses from area 46 and 8 in the frontal lobe, and areas 7a and LIP in parietal cortex were not substantially different and are pooled together for further analyses.

**Figure 3.**
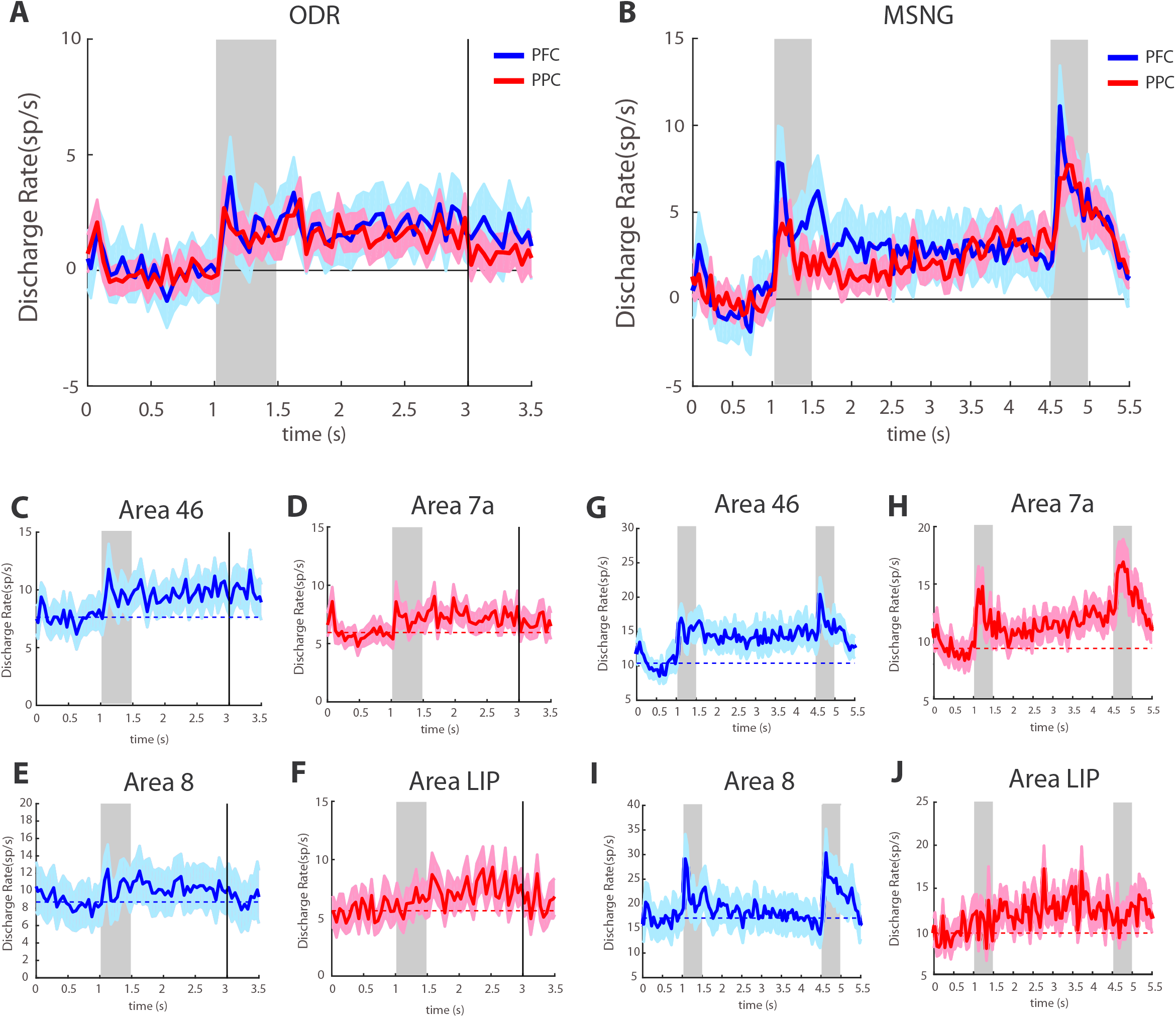
Neural activity in working memory tasks. (A) Average, population peri-stimulus time histogram in the ODR task, for the best available cue location of each neuron. Neurons that showed significantly elevated responses during the cue presentation or delay period of the MSNG task are included (PFC n = 144; PPC n = 145). Gray bar indicates the presence of visual cue in the ODR task. Baseline firing rate, computed in the 1 s fixation period has been subtracted from each histogram, firing rate is plotted relative to this baseline. (B) Average, population peristimulus time histogram in MNSG task for the same neurons as in A. Gray bars represent the presentation of the two visual stimuli in the MSNG task. Firing rate is plotted relative to this baseline. (C-F) Average, population peri-stimulus time histogram for neurons in ODR task, plotted separately for area 46 (n = 96) and area 8 (n = 48) in the PFC and from area 7a (n=97) and area LIP (n=27) in the PPC. Absolute firing rate is plotted here, without subtracting the baseline firing rates. (G-J) Average, population peri-stimulus time histogram for neurons in MSNG task, from the same neurons as in C-F.

### Neuronal responses in correct and error trials

We wished to examine if deviations of firing rate during the delay period of the task in the two areas were predictive of categorical errors about the remembered location of the stimulus. The rationale for our analysis is illustrated in Fig. 4. Activity in the population of neurons that determines working memory behavior can be envisioned as a bump (peak) of activity centered in those neurons that are most activated by the stimulus. Simulation of such a network of recurrent connections revealed that the network behaves as a continuous attractor maintaining the position of the stimulus during the absence of the visual cue (Constantinidis and Klingberg, 2016). This bump is expected to drift randomly during the delay period (Wimmer et al., 2014). The position of the bump at the end of the delay period will determine the location that the subject is actually recalling, and if it is judged to differ from the location of the second stimulus or not (Fig. 4A). In a brain area that does not directly influence what the subject recalls, the drift of such activity would be relatively uncorrelated, however, with the behavioral outcome of the trial.

**Figure 4.**
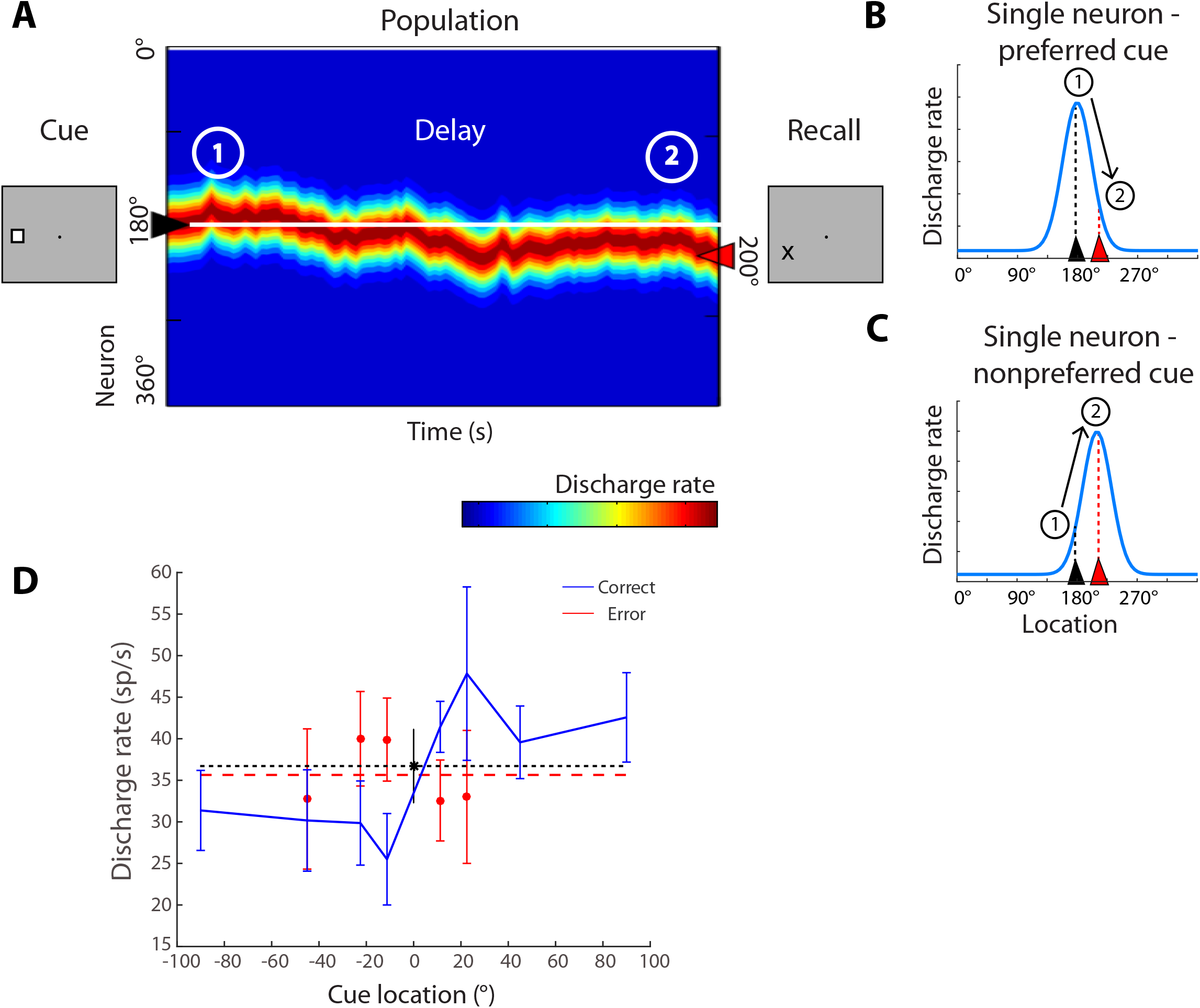
Model of neural activity during the working memory task. (A) Schematic depiction of population neural activity in the neural network representing spatial working memory. Neurons with peak responses at different stimulus locations (indicated as 0 to 360 degrees) are arranged along the y axis of the color plot. The x axis represents time. When the cue appears at the left to the fixation point in the screen (location 180), neurons with peak responses around this location are maximally activated creating a bump of activity in the network. After the cue is no longer present, the population of neurons maintains persistent discharges, however, the bump drifts during the delay period, due to random noise. At the end of the delay period, the location that the subject remembers is determined by the location of the bump in the network, shown here to have drifted counterclockwise (towards the 200 degree location). If a second (nonmatch) stimulus in the MSNG were to appear at that location, the subject would be expected to incorrectly report it as a match. (B) Schematic depiction of changes in neuronal activity at the level of a single neuron with peak response at 180 degrees is shown, for the same sequence of events represented in panel A. At time point 1, shortly after the cue appearance, the neuron is activated maximally as the bump of population responses is centered at 180 degrees. At time point 2, the bump has drifted towards 200 degrees. The neuron’s level of activity, described by its tuning function, is expected to decline. In other words, a neuron is expected to exhibit lower firing rate in error trials than in correct, when the cue appeared at the peak of its tuning function and a nonmatch stimulus followed at its tail. (C) Schematic depiction of changes in firing rate for a different neuron, with a tuning function peak at 200 degrees. At time point A, this neuron is activated moderately. However, as the bump of the activity drifts towards its peak, its firing rate would be expected to increase. Therefore, a neuron is expected to exhibit higher firing rate in error trails than in correct when the cue appeared at the peak of its tuning function and a nonmatch stimulus followed at its tail. D. Firing rate of a single PFC neuron in the delay period involving appearance of the cue at different locations, indicated in the x-axis. The second, nonmatch, stimulus always appears at location 0. Mean firing rate in correct trials (and s.e.m.) is indicated by the blue line; rate in error trials is indicated by the red line. Firing rate in correct trials with the cue at the 0 location (which were followed by a match stimulus) is indicated by the black point; a horizontal line through it is plotted for easy comparison with other locations. Mean firing rate across all nonmatch error trials are indicated by the dashed line.

The model allows us to test predictions at the level of single neurons. If the first stimulus activates a neuron at the peak of its tuning curve, then drifts of the bump that lead into errors will be associated with lower levels of activity for this neuron (Fig. 4B). Therefore, if a cue stimulus appeared at the peak of the neuron’s tuning curve (180° in Fig. 4B), a second stimulus at the tail of the tuning curve (200° in Fig. 4B), and the monkey incorrectly judged this second stimulus as a match, we would expect that the neuron’s activity levels would be lower relative to correct trials. For a neuron for which the original stimulus appeared at the tail of its tuning curve, followed by the second stimulus at the peak (Fig. 4C), we would expect that the neurons’ activity levels would be higher when the subject incorrectly judged the stimulus as a match, compared to correct trials. Importantly, these predictions hold only for neurons in a cortical area that maintains the working memory trace that is read out to determine the categorical judgment. We thus compared the responses of prefrontal and parietal neurons in correct and error trials representing such conditions.

An important point for the analysis is that the model predictions do not depend on the shape of the turning curve, nor do they require it to be Gaussian-shaped over the tested range, such as those depicted schematically in Fig. 4B-C. A real-example tuning curve is shown in Fig. 4D. The reference location is plotted at 0. Delay period firing rate is plotted in the ordinate, following appearance of the cue at each location indicated in the abscissa. The response of the first stimulus at the reference location is indicated by the black point (derived from match trials). We analyzed separately conditions for which the location of the second stimulus corresponded to a lower firing rate than the location of the first stimulus (those above the dotted line, which we refer as “preferred cues”, for simplicity). We also identified conditions for which the location of the second stimulus corresponded to a higher firing rate than the location of the cue stimulus (those below the black dotted line, which we refer to as “non-preferred cues”).

Across the population of prefrontal neurons, firing rate in correct trials involving preferred cues was higher than in error trials, as the model predicted (Fig. 4B). This was also the case for the example neuron (Fig. 4D). A total of 111 PFC neurons with persistent activity in the delay period and error trials follow appearance of the cue at preferred locations in each neuron’s receptive field were available for this analysis. Not all conditions were available for all neurons, so the number of observations varied slightly for each comparison. Average firing rates during the last second of the delay period were 14.5 spikes/s in correct trials and 10.3 spikes/s in error trials (paired *t*-test, *t*_110_ = 4.59, *P* = 1.20 x 10^−5^), as indicated in Fig. 5A.

**Figure 5.**
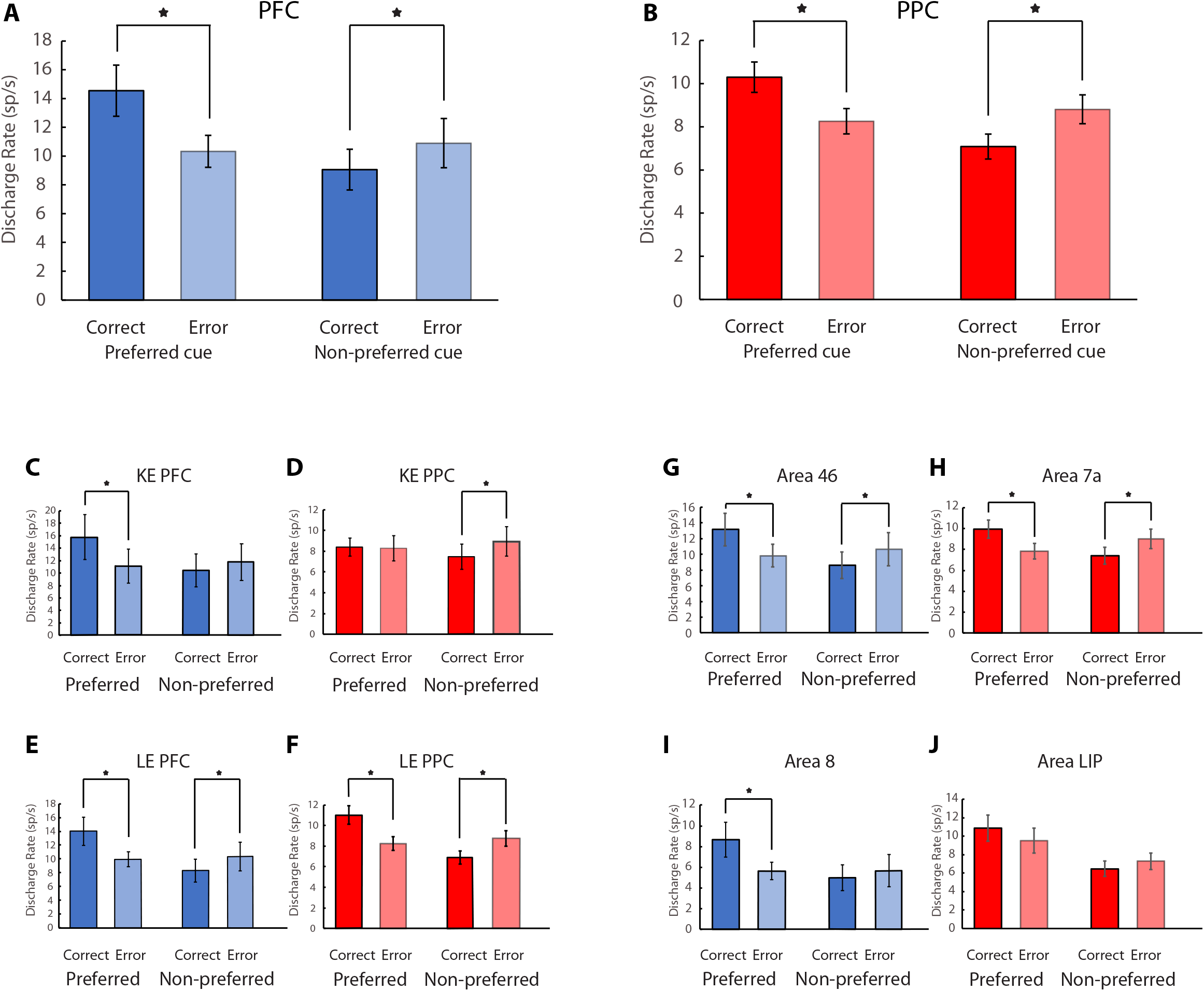
Mean neural activity in correct and error trials. (A) Mean and SEM of firing rates in the last 1s of the delay period for correct and error Nonmatch trials among PFC neurons responsive to the task (n =144). Responses are shown for nonmatch trials involving the cue stimulus presented at a location eliciting higher activity than the second stimulus for each neuron analyzed, indicated as preferred cue (first two bars). Correct and error trials are plotted separately and compared. Star indicates significant difference based on a paired t-test, evaluated at the 0.05 significance level. Responses are also shown for nonmatch trials involving the cue stimulus presented at a location eliciting lower activity than the second stimulus for each neuron analyzed, indicated as non-preferred cue (second two bars). (B) As in A, for PPC neurons (n = 145). (C-F) Results as in A and B, averaged separately for the two monkeys (KE PFC, n = 49; KE PPC, n =44; LE PFC, n =95; LE PPC, n = 101). (G-J) As in A and B, averaged separately for different brain regions (area 46, n =96; area 7a, n = 97; area 8, n = 48; area LIP, n = 27).

Still, this was not a particularly strong prediction of the model, as error trials are likely to include some lapses, and it is well known that error trials typically elicit lower firing rates in the delay period of working memory tasks (Funahashi et al., 1989; Zhou et al., 2013). Most importantly, errors in which the cue appeared at non-preferred locations elicited higher firing rates than correct trials, as predicted by the model (Fig. 4C). This can also be seen in the example neuron (Fig. 4D). A total of 108 PFC neurons with persistent activity in the delay period and error trials following appearance of non-preferred cues were available for this analysis. The average responses in the last second of the delay period were 9.1 spikes/s in correct trials and 10.9 spikes/ in error trials (pair *t*-test, *t*_107_ = −3.26, *P* = 1.49 x 10^−3^). These results confirmed our expectation that the activity of prefrontal neurons during the delay period of the task determines what the subject recalls at the end of the trial (Fig. 5A). It was notable that the mean firing rate across all conditions that the subject reported the second stimulus to be a match (red dashed line in Fig. 4D) were almost identical to the true firing rate that the neuron elicited when the cue stimulus truly appeared in the match, reference location (black dotted line in Fig. 4D).

The result was consistent across subjects, with similar patterns observed for the two monkeys’ prefrontal responses (Fig. 5C, E) and in the two prefrontal areas sampled (Fig. 5G, I), though not all comparisons reached statistical significance. For the preferred cue condition, average responses in the last second of the delay period were greater in correct than error trials in both animals (paired *t*-test, *t*_33_ = 3.35, *P* = 0.002; and *t*_76_ = 3.71, *P* = 0.0004 for the two monkeys, respectively), and in both cortical areas (*t*_73_ = 3.68, *P* = 0.0004 for area 46; *t*_37_ = 2.99, *P* = 0.0049 for area 8). For the non-preferred cue condition, activity in the last second of the delay period was greater in errors than in correct trials for both animals (paired *t*-test, *t*_41_ = −1.59, *P* = 0.12; and *t*_65_ = −2.99, *P* = 0.0039, for the two monkeys respectively) and in both areas (*t*_72_ = −2.94, *P* = 0.0045 for area 46; *t*_35_ = −1.57, *P* = 0.12 for area 8).

Essentially identical patterns of activity were also evident on the population responses of the posterior parietal cortex. Significantly higher responses were observed for correct trials when preferred than non-preferred cues (10.3 vs 8.3 spikes/s, which constituted a significant difference, paired *t*-test, *t*_116_ = 5.46, *P* = 2.75 x 10^−7^). Similarly, the average activity from the posterior parietal cortex for correct trials with non-preferred cues were significantly lower than for error trials (7.1 spikes/s vs. 8.8 spikes/s, paired *t*-test, *t*_114_ = −6.04, *P* = 1.98x 10^−8^; Fig. 5B). We note that the difference between error and correct trials was detectable in the PPC, despite the lower overall firing rate in parietal than prefrontal neurons (Fig. 3B).

In this case, too, results were consistent between subjects (Fig. 5D, F), and in the two areas of the posterior parietal cortex (Fig. 5H, J). For the preferred-cue condition, average responses in the last second of the delay period were greater in correct than error trials, in both animals (paired *t*-test, *t*_29_ = 0.66, *P* = 0.51; and *t*_86_ = 5.90, *P* = 6.95 x 10^−8^, respectively), and in both cortical areas (*t*_73_ = 4.68, *P* = 1.31 x 10^−5^ for area 7a; *t*_23_ = 1.75, *P* = 0.093 for area LIP). For the non-preferred cue condition, activity in the last second of the delay period was greater in errors than in correct trials for both animals (paired *t*-test, *t*_37_ = −3.40, *P* = 0.0016; and *t*_76_ = −4.96, *P* = 4.19 x 10^−6^, respectively) and in both areas (*t*_74_ = −4.70, *P* = 1.18 x 10^−5^ for area 7a; *t*_19_ = −1.18, *P* = 0.25 for area LIP).

Since this analysis represented averages across all neurons that may obscure the patterns of individual neurons, we further examined the phenomenon on a neuron-by-neuron basis, by plotting the neuronal firing rate we observed in each condition (Fig. 6). Each data point in Fig. 6 represents the neuronal average firing rate during the last 1s delay period in correct and error trials, in one condition. For preferred-cue trials, a total of 79% (88/111) PFC neurons exhibited higher firing rate in trials that the monkey correctly identified the second stimulus as a nonmatch, compared to the firing rate in which the monkey erroneously identified the second stimulus as a match (Fig. 6A). This proportion of neurons deviated significantly from a uniform distribution (χ^2^ test, p=6.85×10^−10^). Among PPC neurons, a total of 83% (97/117) exhibited the same pattern of firing rates (Fig. 6C), which was also significantly different than a uniform distribution (χ^2^ test, p=1.09×10^−12^). Conversely, when the cue appeared at a non-preferred location a total of 64% (69/108) PFC neurons exhibited lower firing rate for correct than for error trials (χ^2^ test, p=3.89×10^−3^) in Fig. 6B as did 72% (83/115) PPC neurons (χ^2^ test, p=1.98×10^−6^) in Fig. 6D. We conclude that the effect is highly consistent across neurons in both areas.

**Figure 6.**
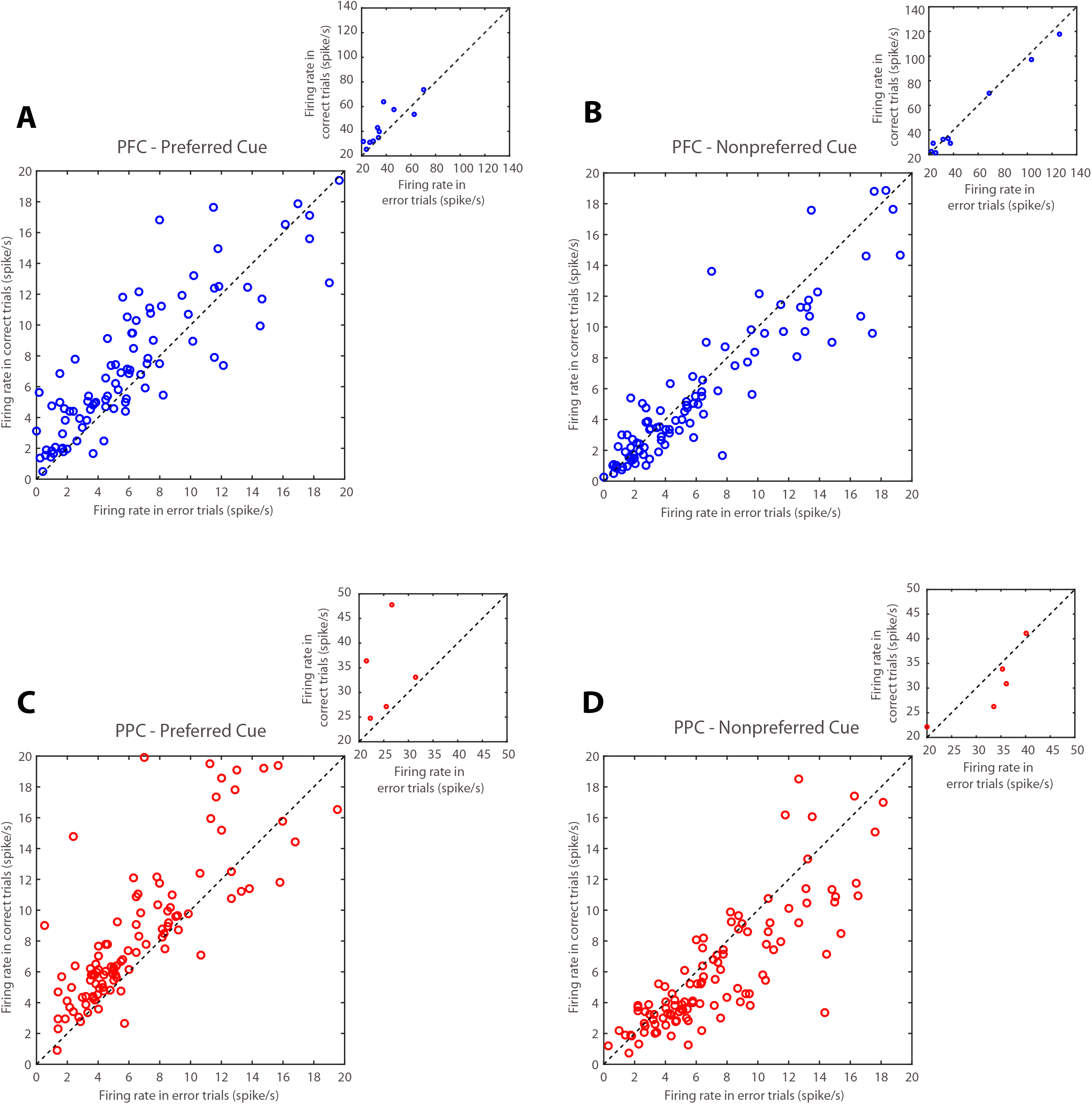
Distribution of neuronal activity in correct and error trials. (A) Neuronal activity from PFC in the last 1s of the delay period from correct and error trials. The data averaged in Fig. 5A, are now plotted individually for each neuron. Each point represents the activity of one neuron in correct and error nonmatch trials for preferred cue trials (n=144). Neurons with firing rate up to 20 spikes/s are represented in the main plot. Outliers, with firing rates >20 spikes/s are shown in the inset. (B) PFC activity in correct and error nonmatch trials from the non-preferred cue condition. (C-D) As in A-B, for PPC neurons (n=145).

### Choice probability analysis

We assessed the reliability with which firing rates can predict the subject’s choice in the MSNG task by performing an ROC analysis that compared the distributions of firing rate in correct and error trials, yielding a quantity also known as choice probability (Britten et al., 1996b). The area under the ROC curve generated by firing rates from correct and error trials represents the choice probability for each neuron. This analysis was performed separately for preferred and non-preferred cues, averaging choice-probability values across neurons in a time-resolved fashion, in successive 500-ms windows, sliding in 50-ms time intervals (Fig. 7).

**Figure 7.**
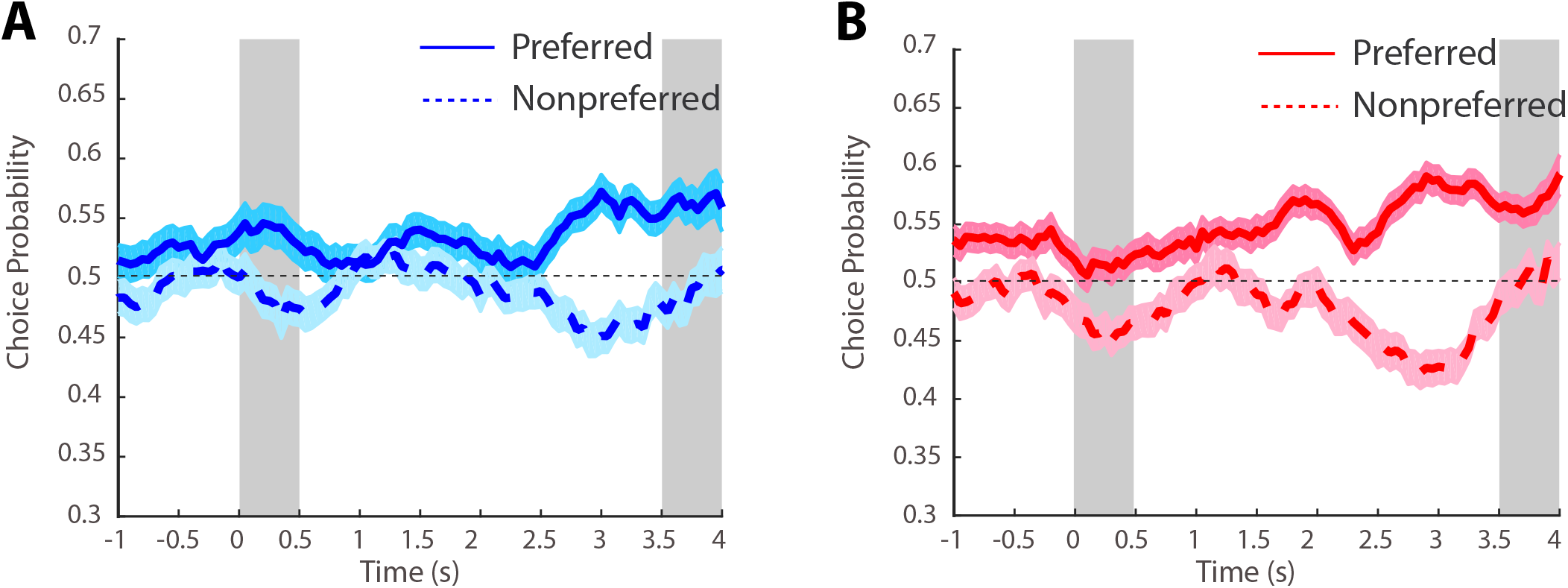
Choice probability. (A) Averaged choice probability from PFC neurons computed in a time resolved fashion and plotted as a function of time across the trial (n=89 for preferred cue; n=88 for non-preferred cue condition). Solid line represents ROC value comparing the distribution of correct and error nonmatch trials, from the preferred cue condition; shaded area around it represents SEM. Dotted line represents ROC value comparing the distribution of correct and error nonmatch trials, from the non-preferred cue condition (B) As in A, data are shown as averaged choice probability from PPC neurons in different conditions (n=91 for preferred cue; n=86 for non-preferred cue condition).

In both PFC and PPC, choice probabilities computed for the preferred cue condition were generally greater than 0.5 (representing chance) through the entire trial (solid lines in Fig. 7). Choice probabilities computed for the non-preferred cue condition were consistently less than 0.5 (dotted lines in Fig. 7). The distance from 0.5 was maximal during the end of the delay period, as would be expected if the animal’s judgement was informed by the firing rate of the neurons at the end of the delay period. Choice probability for the preferred cue conditions computed in the last 1s of the delay period were significantly higher than 0.5 in both areas (two-tailed t-test, *t_88_* = 5.05, *P*=2.40 x 10^−6^ for PFC; *t_90_*= 8.17, *P* = 1.86 x 10^−12^ for PPC). Choice probability for the non-preferred cue conditions computed in the last 1s of the delay period were significantly lower than 0.5 in both areas (two-tailed t-test, *t_87_*= −3.53, *P*=6.75 x 10^−4^ for PFC; *t_85_*= −5.79, *P*= 1.15 x 10^−7^ for PPC). Some subtle differences were present between areas. The probability computed in the 0.5s cue period was significantly different from 0.5 for the preferred cue for the PFC (two tailed *t*-test; *t_88_* = 2.62, *P* = 0.01) and for the non-preferred cue for the PPC (two tailed *t*-test; *t_85_* = −3.43, *P* = 9.39 x 10^−4^). We have previously described PPC activity prior to the appearance of a stimulus, in the baseline fixation interval (of a different working memory task) can be better predictive of behavioral performance, akin to a bias signal (Katsuki et al., 2014). Indeed, we found that the choice probability for the preferred cue condition computed in the fixation period was significantly higher than 0.5 in PPC (two-tailed t-test, *t_90_* = 3.59, P =5.35 x 10^−4^), which is consistent with our previous finding. In sum, these results confirm that the delay period activity in both PFC and PPC was predictive of working memory judgements.

## DISCUSSION

Persistent activity generated during the delay period of working memory tasks represents the properties of remembered stimuli and can account for behavioral fluctuations from trial to trial (Constantinidis et al., 2001; Wimmer et al., 2014). Evidence linking behavior with levels of persistent activity drawn from the Oculomotor Delayed Response task has been criticized, however, on the grounds that may represent motor preparation rather than working memory per se (Lundqvist et al., 2018; Miller et al., 2018). The goal of the present experiments was to determine if persistent activity during a spatial working memory task that required a categorical judgment about two stimuli was predictive of the subject’s response, and if such a relationship was exclusive to the prefrontal cortex or also present in the posterior parietal cortex. We designed a novel task, the Match-Go-Nonmatch-Stay task, which requires a categorical decision based on the remembered spatial location of a stimulus and, arguably, captures better the essence of working memory than delayed response tasks. We often rely on our working memory for categorical decisions and it follows that the neural correlates of working memory should co-vary with the outcome of such judgements. We characterized each neuron’s tuning function and made predictions depending on the relative positions of two stimuli and the subject’s judgment about the remembered location of the first compared to the second. Our results confirmed that trial-to-trial deviations of prefrontal persistent activity were predictive of the subject’s remembered location. Importantly, our task dissociated the response from the spatial location of the remembered stimulus. Thus, our study establishes a direct link between the contents of working memory and persistent activity of prefrontal neurons. Additionally, and contrary to our original expectation, we found that posterior parietal neurons are no less predictive of the subject’s remembered location than prefrontal neurons.

### Neural correlates of working memory

Neurons in the lateral prefrontal cortex and other brain areas that generate persistent activity during working memory tasks that is selective for the properties of the remembered stimuli (Fuster and Alexander, 1971; Kubota and Niki, 1971; Funahashi et al., 1989; Constantinidis et al., 2001). Stimulus location and identity are represented in persistent discharges, as well as more abstract qualities relating to the rules of the working memory task being executed, quantities of stimuli, and categorical judgments, to name a few (Freedman et al., 2001; Crowe et al., 2013; Blackman et al., 2016). Information represented in persistent activity changes based on task demands (Li et al., 2020).Working memory is not the only cognitive domain that persistent neural activity predicts (Constantinidis and Luna, 2019). On a neuron by neuron basis, the ability of a neuron to generate persistent activity in working memory tasks is also predictive of activity generated in other types of tasks, most notably response inhibition (Zhou et al., 2016a). Computational models postulate that persistent activity is sustained by virtue of recurrent connections between neurons with similar tuning for stimulus properties, thus allowing activation to be maintained past the presence of the afferent input resulting in a system that behaves as a continuous attractor (Compte et al., 2000; Wang, 2001; Murray et al., 2017). Structured excitatory and inhibitory connections are both important in the maintenance of working memory, in this scheme (Constantinidis et al., 2002; Wang et al., 2004).

Alternative models that do not rely on persistent activity have been proposed in recent years, relying on activity-silent mechanisms, or rhythmic bursts of discharges (Stokes, 2015; Mi et al., 2017; Lundqvist et al., 2018). A comprehensive discussion of the arguments in favor of and against these models can be found elsewhere (Lundqvist et al., 2016; Riley and Constantinidis, 2016; Constantinidis et al., 2018; Miller et al., 2018). We focus here on one aspect of this debate, the relationship between persistent activity and behavior. Perhaps the strongest argument in favor of persistent activity as the neural correlate of working memory is that discharge rates during the delay period of working memory tasks predict what the subject will recall (Constantinidis et al., 2001; Wimmer et al., 2014), with a level of precession that has not been nearly achieved by alternative models, at least yet. For example, deviations in the discharges of prefrontal neurons have been shown to predict the endpoint of the saccade in the ODR task (Wimmer et al., 2014). Persistent activity recorded from trials in which monkeys made eye movements deviating clockwise vs. counterclockwise relative to the true location of the stimulus yields slightly different tuning curves, as would be expected if the location recalled was determined by the peak of activity at the end of the delay period. Nonetheless, this interpretation was criticized on the grounds that activity may reflect motor preparation to some extent rather then spatial working memory *per se* (Lundqvist et al., 2018). Our current results extend these findings and demonstrate that categorical judgements rather the preparation of a motor movement are influenced by small deviations in persistent activity, in the direction predicted by the bump attractor model.

### Prefrontal and parietal specialization in a bump attractor model

The posterior parietal cortex is a major cortical afferent of the dorsolateral prefrontal cortex (Constantinidis and Procyk, 2004). Posterior parietal and dorsolateral prefrontal cortex share many functional properties with respect to spatial working memory (Rawley and Constantinidis, 2009). Neurons in posterior parietal cortex also generate persistent activity (Gnadt and Andersen, 1988), and this has been shown to represent the remembered locations of visual stimuli, independent of a planned motor response (Constantinidis and Steinmetz, 1996). Tested with the ODR task, virtually identical percentages of neurons exhibiting working memory responses were observed in posterior parietal and dorsolateral prefrontal areas (Chafee and Goldman-Rakic, 1998).

Despite the overall similarity of responses, the prefrontal cortex exhibits some unique properties. For example PPC neurons represent the most recent stimulus location and are disrupted by distracting stimuli (Constantinidis and Steinmetz, 1996; Qi et al., 2010), whereas prefrontal neurons are better able to represent the location of the original stimulus held in memory even after the appearance of distractors, across tasks (di Pellegrino and Wise, 1993; Qi et al., 2010; Suzuki and Gottlieb, 2013), although the specific patterns of responses in the two areas appear to be task-dependent tasks (Jacob and Nieder, 2014; Qi et al., 2015). Persistent activity in prefrontal cortex appears more robust, overall (Masse et al., 2017) and exhibits lower levels of variability in the same tasks (Qi and Constantinidis, 2015). Several studies have thus identified distinct patterns of responses and corresponding roles played by PFC and PPC in working memory and other cognitive functions (Swaminathan and Freedman, 2012; Crowe et al., 2013; Ibos et al., 2013; Jacob and Nieder, 2014; Qi et al., 2015).

Based on these results, one might hypothesize that working memory behavior would depend on the readout of the prefrontal activity exclusively, or more strongly, compared to activity in the posterior parietal cortex. Contrary to this expectation, we found that PPC the activity of prefrontal neurons was no less predictive of behavioral outcomes than prefrontal activity. Two explanations could account for these results. First, prefrontal and posterior parietal cortex simultaneously, and independently of each other influence downstream areas that ultimately determine behavior based on recall of information maintained in memory. Under such a model, “readout” of memory activity occurs in both areas. Alternatively, prefrontal cortex is the ultimate arbiter of information held in memory, however posterior parietal activity determines prefrontal activity to a large extent. Prefrontal and parietal neurons are co-active in working memory tasks, with variations in parietal cortex preceding in time of similar changes in prefrontal cortex (Crowe et al., 2013). Under the second scenario, our results suggest precise propagation of deviations observed in the PPC into the PFC. Future experiments could differentiate between these alternatives.

## ACKNOWLEDGMENTS

Research reported in this paper was supported by the National Eye Institute and National Institute of Mental Health of the National Institutes of Health under award number R01 EY017077 and MH116675. We wish to acknowledge Kathini Palaninathan, Austin Lodish, Du Gu, and Leonardo Silenzi for technical help; Rob Hampson, Emilio Salinas, Benjamin Rowland, and Wenhao Dang for helpful comments on the manuscript.

